# Global motor system suppression as the primary mechanism of human action stopping: challenging the pause-then-cancel model

**DOI:** 10.1101/2025.05.13.653703

**Authors:** Eashan Ray Chaudhuri, Tiombe Long, Ricci Hannah

## Abstract

The ability to stop a planned or ongoing action is fundamental to inhibitory control. A recent theory proposes that stopping involves two distinct phases: an initial global suppression of motor activity (“pause”) followed by a selective cancellation of the targeted action. However, the necessity of a second “cancel” stage remains debated. We tested whether global suppression alone is sufficient to stop movement by analysing electromyography from task-relevant agonist and antagonist muscles, alongside transcranial magnetic stimulation measures of global motor suppression from task-irrelevant muscles, during a stop-signal task in adult human participants of both sexes. In Experiment 1, reanalysis of ballistic finger movements revealed that agonist muscle offset consistently preceded behavioural stopping, aligning with the time course of global suppression. In Experiment 2, we extended these findings to whole-arm reaching movements, demonstrating that global motor suppression persisted beyond the termination of muscle activity when stopping prevented movement initiation, but disengaged in time for antagonist activation used to interrupt movements once they had begun. These findings challenge the pause-then-cancel model, instead supporting a single-stage global suppression framework. They also suggest that the global suppression mechanism is not a rigid, top-down stopping mechanism but rather part of a broader motor control system that flexibly adjusts movement commands based on task demands.

**Significance Statement:** A recent theory suggests that stopping an action involves a two-stage process: first pausing movement execution, then selectively cancelling specific motor commands. Our study challenges this view, demonstrating that global motor suppression alone is sufficient to terminate motor commands. Using electromyography and transcranial magnetic stimulation during two distinct motor tasks, we show that suppression fully curtails agonist motor commands before behavioural stopping, without requiring a distinct cancellation step. These findings reshape our understanding of action stopping and provide a foundation for exploring the pathophysiology of movement and neuropsychiatric disorders associated with impaired behavioural suppression.

## INTRODUCTION

Many of us can recall moments when we were about to act but refrained at the last moment, like suppressing an angry outburst or reaching for a pan on the stove before realising it’s hot. This process, known as action stopping, is believed to involve a prefrontal-basal ganglia-thalamocortical network (Jahanshahi et al., 2015; Hannah and Aron, 2021). While evidence supports this network, key questions remain about how it achieves movement suppression.

Traditionally, action stopping has been viewed as a single-stage process where a global suppression of motor activity, evidenced by a reduction in corticospinal excitability measured with transcranial magnetic stimulation (TMS) over task-irrelevant motor cortical representations (Badry et al., 2009; Greenhouse et al., 2012; Jana et al., 2020), is sufficient to terminate movement. This suppression is thought to be mediated by the subthalamic nucleus (STN) (Wessel et al., 2016, 2022), and in this framework, accounts for stopping without requiring additional processes (Hannah and Aron, 2021).

However, a competing proposal, the “pause-then-cancel” model, suggests stopping unfolds in two stages (Schmidt and Berke, 2017): a global “pause” that temporarily suppresses motor activity, followed by a more selective “cancel” process that targets specific movement commands. Here, the global suppression represents only the initial pause stage (Diesburg and Wessel, 2021).

Several human studies have tested predictions of the pause-then-cancel model, with some evidence supporting the involvement of distinct pause and cancel processes (Tatz et al., 2021, 2024; Wadsley et al., 2023; Hervault and Wessel, 2025; Wu et al., 2025). The model itself was originally developed to explain early STN activity observed in rodent studies, where activation begins within ∼15 ms of a stop cue and well before movement cessation (Schmidt et al., 2013). However, this early activation is not mirrored in humans or non-human primates, where STN activity typically emerges closer to the time of behavioural stopping (Bastin et al., 2014; Benis et al., 2016; Pasquereau and Turner, 2017). This discrepancy raises a question about whether such a two-stage process is truly necessary to explain human stopping.

A direct way to evaluate this question is to examine the suppression dynamics on trials where muscle activity is initiated but then interrupted before producing overt movement. Previous work suggests that the initial decline in muscle activity follows the global suppression and precedes the behavioural latency of stopping, the stop signal reaction time (SSRT) (Raud and Huster, 2017; Hannah et al., 2020; Jana et al., 2020; Raud et al., 2022). Interestingly, recent work proposed that the P3 event-related potential, which emerges in scalp EEG around SSRT, might reflect the secondary “cancel” process (Hervault and Wessel, 2025; Hervault et al., 2025). However, if muscle activity is completely suppressed before SSRT, and hence before typical P3 onset, it would suggest that global suppression alone is sufficient to terminate movement, undermining the need for a distinct cancellation stage.

Beyond this theoretical debate, a wider question remains: how does global suppression operate during real-world actions? Most stopping studies focus on simple, ballistic movements, like button presses or saccades (Greenhouse et al., 2012; Jana et al., 2020; Tatz et al., 2021, 2024), which offer little opportunity for interruption once movement has begun. However, many natural actions, such as reaching for a hot pan, unfold over longer durations, allowing for suppression both before and after onset. Such cases require both terminating agonist muscle activity and activating antagonist muscles to brake the movement (Kudo and Ohtsuki, 1998; Atsma et al., 2018). This raises a critical question: does the same global suppression mechanism, well-documented in ballistic tasks, also operate when halting non-ballistic but naturalistic movements that are already underway but not yet complete? If so, how does it avoid disrupting necessary antagonist activity required for braking?

To address these questions, we conducted two experiments investigating the temporal dynamics of motor suppression in relation to agonist and antagonist muscle activity during ballistic button-press and non-ballistic whole-arm reaching versions of the stop signal task.

## MATERIALS AND METHODS

### Participants

Healthy, adult, human volunteers provided written informed consent to participate in two experiments. The experiments were approved by the UCSD Institutional Review Board (protocol #171285) and the Research Ethics Committee at King’s College London (HR/DP-21/22-31662).

Experiment 1. Eighteen participants (11 females; age 19 ± 0.4 years; 15 right-handed) were recruited as part of a separate study (Jana et al., 2020). In this study, we reanalysed a subset of the data and present new findings. One person’s data was excluded for poor behavioural performance.

Experiment 2. Twenty-two participants (7 females; age 23 ± 1 years; 19 right-handed). Two people’s data were excluded for poor quality behavioural and/or EMG data. The sample size was based on our previous work noted above (Jana et al., 2020), which used a similar paradigm and task parameters, and successfully demonstrated robust effects, including the relationship between peak EMG timing and SSRT, as well as condition × time interactions in global MEP suppression (n = 17). This sample size also aligns with recent studies using similar methods to examine global motor suppression (Raud et al., 2020; Wadsley et al., 2023).

### Stop signal task

The tasks were coded using MATLAB (2016b and 2022b; Mathworks, USA) in conjunction with Psychtoolbox (Brainard, 1997). Experiment 1, which has been described in detail elsewhere (Jana et al., 2020), involved participants responding to white arrows on a screen by pressing keys with either their left index or pinky finger, depending on the arrow’s direction. Participants had 1 second to make a response and were encouraged to respond as quickly and accurately as possible. Trials requiring a response, termed “Go” trials, accounted for 75% of all trials. In the remaining 25% of trials, the arrow turned red after a variable delay (the stop signal delay, SSD), signalling participants to try to stop the impending movement. The SSD increased by 50 milliseconds after a Successful Stop and decreased by 50 milliseconds after a Failed Stop.

The task in Experiment 2 (Figure 1A) built upon our recently developed whole-arm reaching version of the stop signal task (Hannah et al., 2022). In this version, participants moved a computer mouse cursor towards targets displayed on the screen. The sensitivity of the computer mouse was intentionally reduced so that participants were required to make larger, whole-arm movements, ∼10 cm displacement in the anterior–posterior plane, to reach the targets. Each trial began with participants positioning the cursor inside a designated small square, referred to as the ‘home pad’. One second after the cursor entered the home pad, two blue target squares and a Go signal (a letter) appeared simultaneously on the screen. The letter ‘Q’ instructed participants to move the cursor toward the left target, while ‘O’ indicated movement toward the right target. These cues were intentionally chosen for their visual similarity, in order to slow response initiation and reduce the likelihood of SSD floor effects or violations of context independence (Bissett et al., 2021). Participants were instructed to execute their movements in a single, smooth motion, aiming for both speed and accuracy. Trials requiring a response were termed Go trials. Failure to reach the target within the allotted time of 1.5 seconds resulted in a trial timeout, accompanied by a message indicating ‘Too slow’.

**Figure 1.**
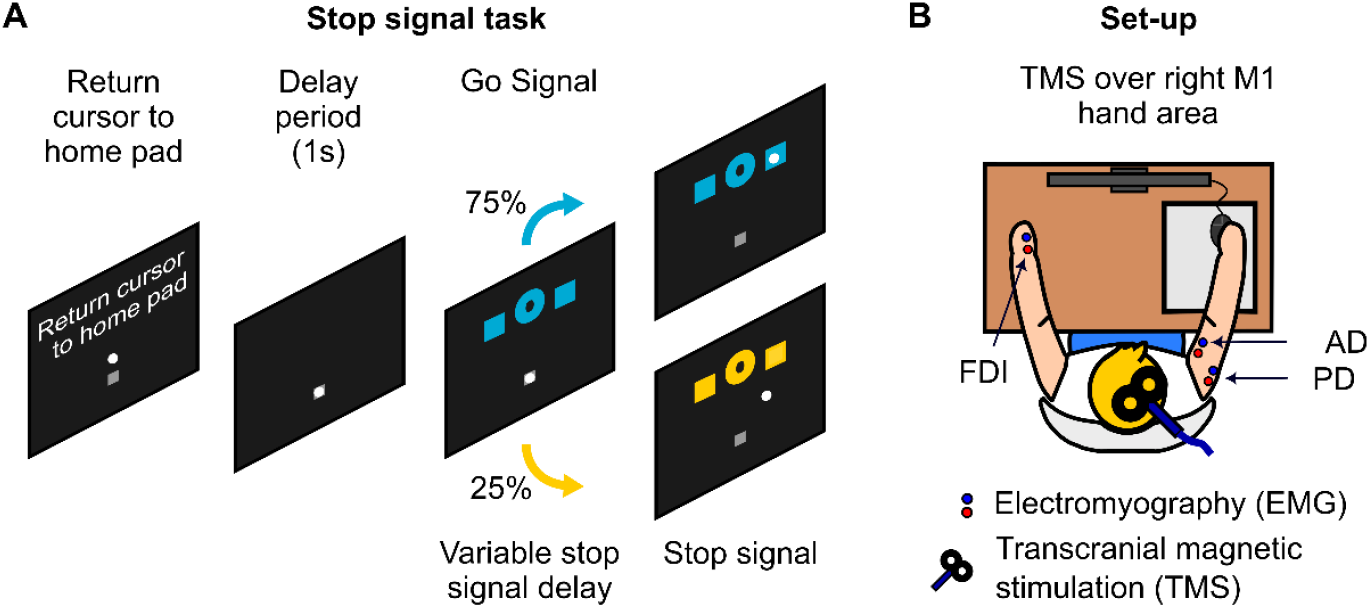
Experimental procedures and set-up employed in Experiment 2. **A**. Sequence of events in the whole-arm reaching stop-signal task. Participants controlled a cursor using a computer mouse, starting each trial in the home pad before receiving a directional cue (‘Q’ for left, ‘O’ for right) to move to one of two target squares. In Go trials (75%), participants made the cued movement. In Stop trials (25%), a stop signal (colour change) instructed participants to cancel the movement after a variable stop-signal delay (SSD). **B**. Schematic of the experimental setup. Participants were seated in front of a screen, with EMG electrodes recording from the right anterior deltoid (AD, agonist), right posterior deltoid (PD, antagonist), and the left first dorsal interosseous (FDI, task-irrelevant muscle). A TMS coil over the right motor cortex elicited motor evoked potentials (MEPs) in the left FDI to assess global motor suppression, with TMS delivered at fixed time points relative to the stop signal or its yoked equivalent on Go trials.

As before, in a subset of trials (25%) a stop signal, indicated by a colour change, was introduced following a variable SSD. Here, the SSD increased by ∼33.3 milliseconds after a Successful Stop and decreased by ∼33.3 milliseconds after a Failed Stop (Hannah et al., 2022). Participants were instructed to halt their movements promptly upon presentation of the stop signal. For the purposes of adaptive tracking, any movement that caused the mouse cursor to exit the home pad was classified as a response (i.e., a failed stop), whereas successful stopping was defined as the cursor remaining within the home pad.

However, in practice, participants were almost always able to interrupt the ongoing movement before it reached the target [∼96% stop trials; and see (Hannah et al., 2022)]. This allowed us to study stopping at different points during movement execution, prior to, just after, and well after initiation. While this operational definition was useful for adaptive tracking, many of these so-called failed stops were in fact successful interruptions of action.

We therefore refer to these trials in Experiment 2 as “Early Stops” and “Late Stops” when discussing the data, to reflect the timing of inhibition within the movement rather than to imply failure or success in the conventional stop-signal sense. Importantly, although Late Stop trials involved partial movement, they were considered instances of successful stopping, as the action was cancelled prior to reaching the target. Complete failures to reach interrupt the movement before the target were termed “trigger failures”, assumed to represent failures to initiate the stop process altogether (Hannah et al., 2022).

At each block’s end, participants received feedback on their average total response time. They were encouraged to speed their responses if they began to slow down or to exert their best effort to stop if stopping accuracy fell below 30%. Participants rested for as long as they needed between blocks.

### Electromyography (EMG)

In Experiment 1, surface EMG signals were recorded from the left first dorsal interosseus (FDI) and abductor digiti minimi (ADM) muscles. Signals were band-pass amplified (×5000; Grass QP511 AC amplifier, Grass Instruments, USA) with a cut-off frequency ranging 30 and 1000 Hz. Signals were then digitised at 1000 Hz (Micro 1401 mk II, Cambridge Electronic Design, Cambridge, UK) and recorded via data acquisition software (Signal version 4, Cambridge Electronic Design, Cambridge, UK). Note that EMG was also recorded from the task-irrelevant extensor carpi radialis muscle, to record motor evoked potentials (MEPs) during the task. However, our focus here is on the task-relevant FDI and ADM muscles.

In Experiment 2, surface EMG was recorded from the anterior and posterior deltoid (AD and PD) muscles of the task-relevant right arm (Figure 1B). These muscles served as agonists and antagonists in the reaching movement. EMG signals were obtained using a bipolar arrangement over the muscle, with a ground electrode situated over the acromion. Surface EMG was additionally recorded from the task-irrelevant FDI muscle of the left hand (Figure 1B), which was not directly involved in the task. In each trial, signals were captured from 1 second preceding the Go cue to 1.5 seconds following it.

EMG signals underwent band-pass amplification (×1000; D360, Digitimer, UK) with a frequency cut-off ranging from 10 to 2000 Hz. Sampling occurred at a rate of 4 kHz using the Power 1401 mk II device (Cambridge Electronic Design Ltd., UK). Data was acquired via Signal v8 software (Cambridge Electronic Design Limited, UK).

### Transcranial magnetic stimulation (TMS)

In Experiment 2, MEPs were evoked using a TMS device (Magstim 200^2^, The Magstim Company Ltd., UK) delivering monophasic pulses, and connected to a figure-of-eight coil (70 mm diameter, The Magstim Company Ltd., UK). The coil was positioned on the scalp over the right primary motor cortex (M1) representation of the left FDI muscle and oriented so that the coil handle was approximately perpendicular to the central sulcus, that is at ∼45° to the mid-sagittal line, and the initial phase of current induced in the brain was posterior-to-anterior across the central sulcus (Figure 1B).

Prior to the experiments, the motor hot spot was determined as the position on the scalp where slightly supra-threshold stimuli produced the largest and most consistent MEPs in FDI. The position was marked on a cap worn by the participants. We then established the test stimulus intensity to be used during task, which was set to produce a mean MEP amplitude of approximately 1 mV whilst the participant was at rest.

MEPs were also evoked in the right AD muscle before beginning the main experiment to estimate the corticomuscular conduction time to the deltoid muscles. After determination of active motor threshold (AMT), 10 stimuli were delivered at 150% AMT during slight voluntary contraction (∼10% of maximum EMG).

### Procedure

In each experiment, participants began with a brief practice block to familiarise themselves with the task and to estimate an appropriate initial SSD for each individual. This titration was necessary because participants vary in their reaction times and SSRTs, and the staircase algorithm used during the main task requires a starting SSD that is reasonably close to the value that yields ∼50% stopping accuracy.

Both Experiment 1 and 2 involved 12 blocks, each block consisting of 96 trials. During the task of Experiment 2, TMS was administered on both stop trials and on 50% of Go trials. In Experiment 2, on five out of every six Stop trials, a single TMS pulse at the test stimulus intensity was delivered at one of several time points: 100 ms, 150 ms, 200 ms, 250 ms, and 300 ms after the Stop signal. In the remaining one out of six Stop trials, no TMS was delivered. The latter helped minimise the total number of pulses administered. On Go trials, TMS was synchronised with the timing of the Stop signal on the preceding Stop trial. Additionally, TMS was delivered at the time of the Go signal, serving as a baseline estimate of excitability. Consequently, there were 48 trials per TMS time point on Stop trials in Experiment 2.

### Data analysis

All analyses were performed using custom scripts written in MATLAB (2024b; Mathworks, USA).

### Behavioural performance

SSRT in both experiments was determined via the integration method (Verbruggen et al., 2019), which is based on the Race Model (Logan et al., 1984), hence termed SSRT_RM_. The model assumes a race between a go process and a stop process competing for completion. We also obtained SSRT estimates using a kinematic approach by identifying the peak of the cursor velocity-time curve in Late Stop trials as an estimate of when action stopping occurred [SSRT_K_; (Hannah et al., 2022)]. Thus, SSRT_RM_ reflects stopping before detectable movement execution (Early), whereas SSRT_K_ captures the interruption of an ongoing movement (Late).

To derive SSRT_K_, we calculated resultant cursor displacement relative to the home pad, estimated resultant velocity, and measured the time of peak velocity on Late Stop trials relative to the stop signal to derive SSRT_K_. Unlike SSRT_RM_, SSRT_K_ does not rely on the assumptions of the independent Race Model (e.g., context independence or a single stopping process), offering a more direct, movement-based estimate of stopping latency. Its close correspondence with SSRT_RM_ across participants (see Results) supports the validity of both measures as behavioural indices of stopping despite their differing underlying assumptions.

Notably, both SSRT_RM_ and SSRT_K_ were previously validated against the BEESTS algorithm (Hannah et al., 2022), which explicitly models trigger failures (Matzke et al., 2017). SSRT values derived via BEESTS closely matched those from our methods, and inferred trigger failure rates were very low (∼1%). A similar pattern was observed in the current study, with rare trigger failures and tight agreement between SSRT_RM_ and SSRT_K_ (see Results), further supporting the robustness of these behavioural stopping estimates.

In Experiment 2, Go reaction times were estimated as the time from the Go signal to the time when the cursor first left the home pad, whilst total response times encompassed the time from the Go to when the cursor entered the target.

Errors on Go trials in Experiment 2 included omissions, missed targets, incorrect target hits, and false alarms. Kinematic data were also used to identify trials where the initial movement was in the wrong direction. Specifically, these errors were defined as trials where the cursor moved to the incorrect side of the midpoint of the screen for at least six consecutive frames (∼100 ms). These trials were classified as errors to avoid interpretive ambiguity, since suppressive mechanisms may contribute to the subsequent correction, potentially confounding the assessment of Go trials. We also quantified trigger failures, reflecting failure to trigger the stop process, as the proportion of stop trials in which there was a complete failure to interrupt the movement before it reached the target (Hannah et al., 2022).

### EMG processing

EMG analysis followed a similar approach to our previous work (Hannah et al., 2020; Jana et al., 2020), with slight adjustments to the pre-processing. Data from Jana et al. (2020) in Experiment 1 were reanalysed using the updated pre-processing method to ensure consistency across studies.

In all cases, EMG signals were interpolated from -1 to 1.5 ms around the TMS to remove the artefact. The agonist and antagonist signals were then bandpass filtered between 10 and 2000 Hz using a 2^nd^-order Butterworth filter (roll-off 24 dB/octave), and bandstop filtered to remove mains hum (58-62 Hz for Experiment 1 and 48-52 Hz for Experiment 2). After filtering, signals were full-wave rectified, and the root mean square (RMS) was computed with a centred 20 ms window. Task-irrelevant FDI EMG data in Experiment 2 were bandpass filtered as above.

EMG bursts were identified as peaks exceeding 10% (Experiment 1) or 15% (Experiment 2) of the average peak EMG activity during correct Go trials. A higher threshold was applied in Experiment 2 due to the increased variability and background activity associated with the reaching movement, where EMG activity supports both movement and limb stability at the start and end points. From each peak, the onset was determined by backtracking to the point where the signal fell below 5% (Experiment 1) or 10% (Experiment 2) of the peak for at least 20 consecutive milliseconds (Figure 2A and 2B). Similarly, the offset was marked by tracking forward. The onset of EMG activity decline was simply defined as the time of the peak. Outliers in EMG timing were identified as values exceeding 1.5 times the interquartile range (IQR) of the first and third quartiles. Additionally, the algorithm occasionally captured early PD activity characterised by a slow rise beginning shortly after agonist onset (MacKinnon and Rothwell, 2000), rather than later burst-like activity triggered by the stop signal. Therefore, trials where PD onset occurred within 50 ms of stop signal were not included in subsequent analyses.

**Figure 2.**
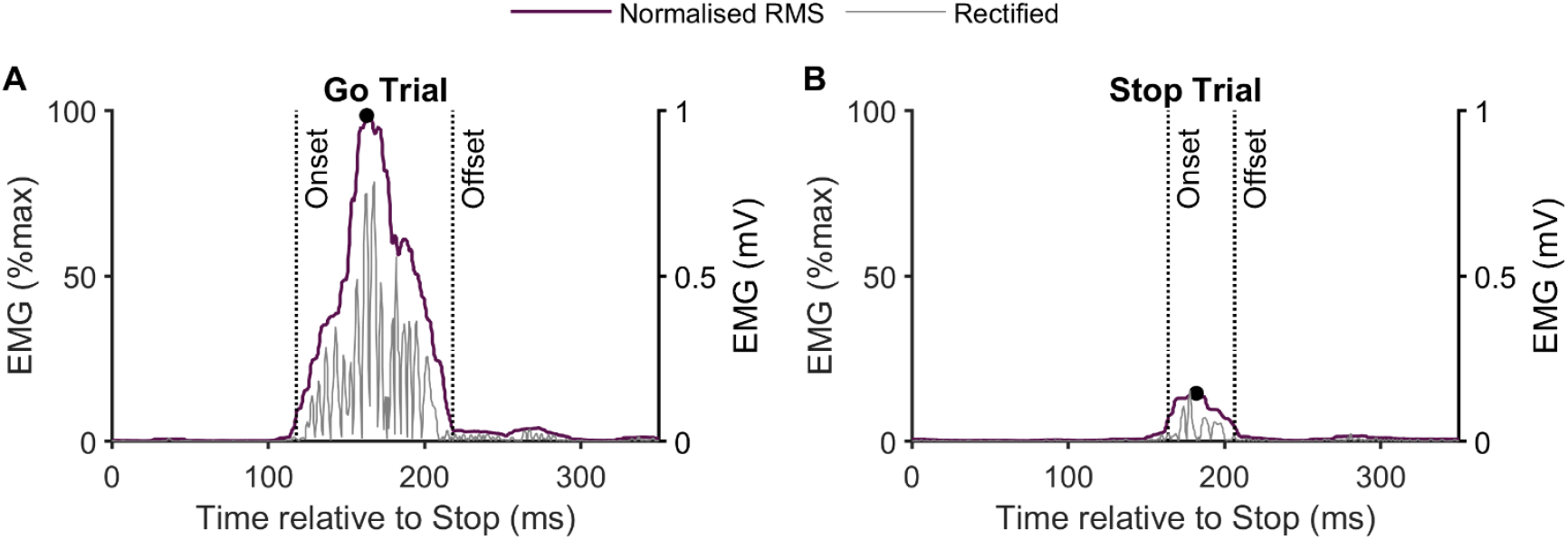
Representative data from a single participant showing normalised RMS EMG and absolute rectified EMG traces from the FDI muscle for single trials, along with the timing of EMG burst onsets, peaks (black circle) and offsets. **A**. Go trial. **B**. Stop trial. The minimal smoothing window applied to the RMS EMG provides reliable estimates of burst onsets, peaks and offsets.

Agonist muscle EMG amplitudes (FDI, Experiment 1; AD, Experiment 2) were normalised to the mean peak agonist activity in Go trials (% max). For the antagonist PD muscle in Experiment 2, EMG amplitudes were normalised to the mean peak PD activity in Late Stop trials (% max).

In Experiment 1, we focused on the timing of both the decline and the offset of EMG activity in the agonist FDI and ADM muscles relative to the stop signal, and the behavioural latency of stopping, SSRT_RM_. Our primary aim was to demonstrate that agonist muscle activity terminated before the SSRT_RM_. Analyses were restricted to successful stop trials that contained suprathreshold EMG bursts. Trials without detectable EMG activity were excluded from these analyses.

Note that while EMG peaks (marking the onset of decline) are present in both go and stop trials, their characteristics differ systematically across conditions. In stop trials, the peak occurs earlier, has a smaller amplitude, and marks the point at which EMG activity begins to decline rapidly, coinciding with the onset of global motor suppression (Jana et al., 2020). This suggests that the peak itself reflects the initiation of inhibition in stop trials and is informative about the timing of stopping.

In Experiment 2, we extended this investigation to the agonist AD and antagonist PD muscles. Specifically, we analysed the timing of EMG decline and offset in the agonist muscle, and the onset of EMG activity in the antagonist muscle, relative to the stop signal, SSRT_RM_ and SSRT_K_ and the global suppression. The goals were to replicate Experiment 1’s findings by confirming a rapid termination in agonist activity before the SSRT, and to explore how this termination corresponded with the timing and duration of global suppression. Additionally, we investigated the relative timing of antagonist muscle onset and global suppression.

For Experiment 2, MEP amplitudes in the task-irrelevant left FDI muscle were measured trial-by-trial within a 30 ms window from 18 ms after the TMS. Data were included if FDI EMG activity in the 100 ms before the TMS was < 0.05 mV. MEP amplitudes recorded at the Go signal in Go trials were averaged as a baseline for comparison across other TMS time points (100, 150, 200, 250, 300 ms after the Stop signal). For each TMS time point, MEP data were averaged within each trial type (Correct Go, Early Stop, Late Stop) and expressed as a ratio relative to Go signal MEP amplitudes.

### Corticomuscular conduction time

The AD MEP onset latency across 10 trials was identified visually, and corticomuscular conduction time was taken as the minimum value across all stimuli (Hamada et al., 2013; Hannah and Rothwell, 2017).

### Statistical analyses

Data are presented as mean ± SEM. Statistical significance for all analyses was set at p < 0.05.

In Experiment 1, agonist EMG timings (peak, offset) were averaged across the FDI and ADM muscles and compared to the SSRT using paired-samples t-tests. The relationship between EMG timings and SSRT were assessed using robust linear regression (‘robustfit’ function, MATLAB). Cohen’s *d* was employed as an estimate of effect size.

In Experiment 2, EMG timings for the agonist (peak, offset) and antagonist (onset) muscles were compared to SSRT using paired-samples t-tests, and their relationships with SSRT were assessed using robust linear regression. A repeated-measures ANOVA was conducted to assess the effects of trial type (Go, Early Stop, Late Stop) and TMS time point (100–300 ms) on task-irrelevant FDI MEP amplitudes. *Post hoc* paired t-tests were conducted to examine significant differences between trial types at each time point. Additional paired t-tests compared MEP amplitudes at each time point and trial type to baseline MEP amplitudes to determine whether they were significantly below baseline. The similarity of SSRT_RM_ and SSRT_K_ were evaluated via paired t-tests and robust linear regression, along with Bayes Factor (BF_10_).

## RESULTS

### Experiment 1

We reanalysed EMG data from a previous study (Jana et al., 2020) to examine the timing of EMG offset in relation to the latency of stopping, SSRT.

As previously reported (Jana et al., 2020), participants exhibited typical stop-signal task performance, with Go RTs of 430 ± 17 ms, Failed Stop RTs of 391 ± 12 ms, mean SSD of 194 ± 18 ms, and pStop of 49 ± 1%. Mean Failed Stop RT was therefore shorter than Go RT at the group level and in 15/17 individuals. Notably, even in Successful Stop trials, small bursts of agonist muscle activity were observed, suggesting that these bursts reflected initiated but subsequently stopped motor commands before they could elicit a behavioural response. The peak of agonist EMG (160 ± 9 ms), marking the transition from rising to falling activity, preceded SSRT (217 ± 6 ms) and was itself preceded by global motor system suppression by an interval corresponding to corticomuscular conduction time. This indicates a direct link between muscle activity suppression and global motor suppression (Jana et al., 2020).

This prompted the current analysis, which aimed to accurately determine the timing of complete agonist (FDI and ADM) EMG suppression by pinpointing the offset time in typical Successful Stop trials. EMG bursts were detected in 70 ± 4% of Successful Stop trials, with smaller amplitudes compared to Go trials, consistent with their cancellation (Figure 3A). The onset of the EMG decline (i.e. peak) occurred 155 ± 8 ms after the stop signal, closely aligning with our previous estimates (Jana et al., 2020). When plotted relative to the stop signal, the EMG bursts in Successful Stop trials appeared to decline gradually, extending beyond SSRT (Figure 3A). This could suggest the need for an additional cancellation process, as proposed by the pause-then-cancel model.

**Figure 3.**
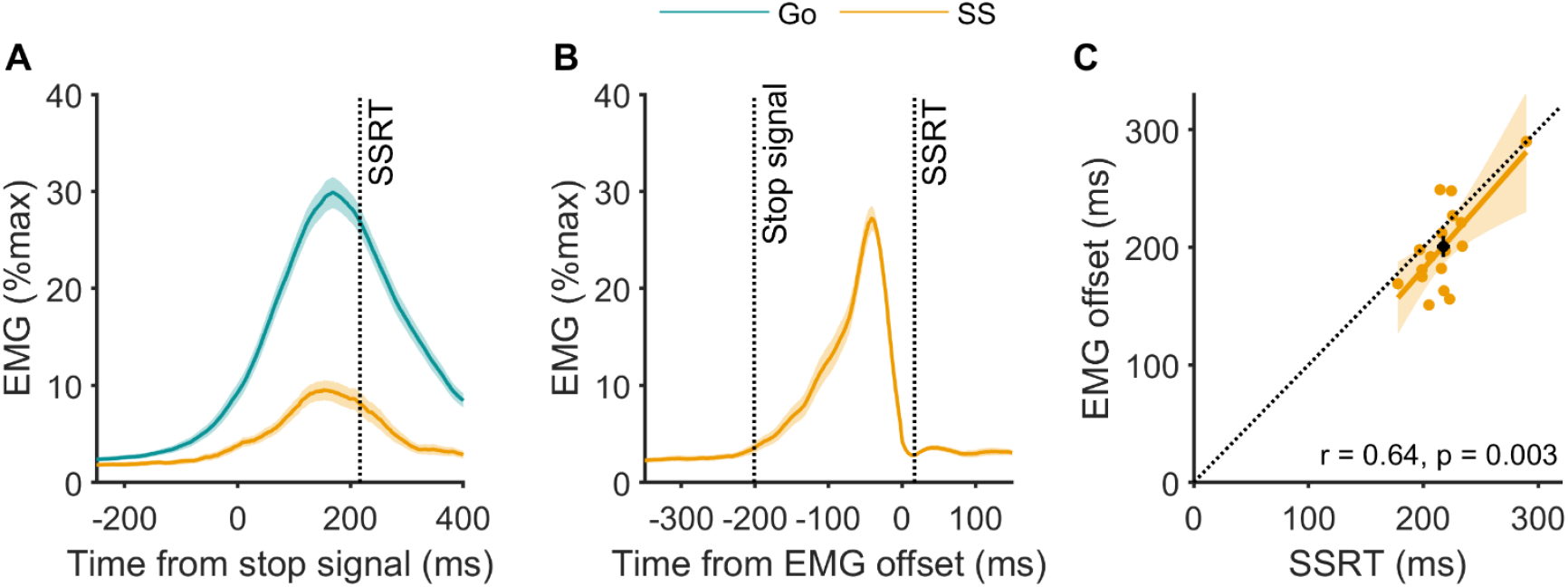
Group average EMG amplitudes from the agonist muscles (FDI and ADM) in Go and Successful Stop trials during Experiment 1. Timings are displayed relative to the stop signal (A) and agonist EMG offset (B), with shaded areas representing the SEM. **A**. Illustrates the lower amplitude and shorter duration of partial EMG bursts on Successful Stop trials, yet gives the impression of a relatively prolonged decline in EMG, seemingly extending beyond the behavioural stopping latency (SSRT_RM_). **B**. Provides a clearer picture of the rapid termination of EMG activity, emphasising the swift decline from the peak to pre-burst levels. **C**. Shows a significant positive relationship between SSRT_RM_ and the time of agonist EMG offset. The dashed black line represents unity, and the black circle marks the group mean and SEM for SSRT and EMG offset. The group mean, along with most individual data points (yellow circles), fall below the unity line, indicating that EMG offset occurred before SSRT for most participants. Note: The apparent extension of EMG bursts beyond SSRT in panel A is due to temporal smearing caused by variability in burst timing across individuals. Aligning to EMG offset (panel B) reduces this effect and provides a clearer picture of muscle suppression dynamics.

However, this apparent persistence is misleading and likely reflects temporal smearing in the group-averaged data. Inter-individual variability in SSRT and EMG burst timing tends to flatten averaged waveforms, particularly when signals are time-locked to external events like the stop signal. This phenomenon has been previously documented in the stop-signal literature (Salomoni et al., 2025), and can obscure the true temporal dynamics of suppression. Aligning to intrinsic physiological events, such as EMG offset, reduces this smearing and provides a more accurate picture of when muscle activity actually terminates.

A clear illustration of temporal smearing can be seen in Figure 3A, where Go trial EMG though normalised to 100% for each participant, reaches only ∼30% in the grand average. This under-representation arises from variability in EMG timing across trials and participants, which flattens the averaged trace when aligned to the stop signal. A similar effect is evident for Successful Stop trials: EMG bursts appear smaller in Figure 3A than in Figure 3B, where the data are aligned to EMG offset. This difference does not reflect a change in signal magnitude, but rather a reduction in temporal variability when aligning to a physiologically meaningful event. Together, these examples demonstrate how time-locking to external events like the stop signal introduces smoothing that can obscure not only the true amplitude and shape of EMG bursts, but also their precise timing relative to other external events, most importantly here, behavioural stopping.

Our new analysis reveals that the agonist EMG burst was fully suppressed (offset) at 201 ± 9 ms after the stop signal, just 46 ± 4 ms after the initial decline. Aligning the offset of EMG bursts across Successful Stop trials reveals the rapid return of EMG to near-baseline levels (Figure 3B). By the time of SSRT, the EMG signal is ∼3% of the peak EMG in Go trials, close to the baseline level of ∼2%, with the absolute difference being negligible compared to the signal’s dynamic range (the greatest peak EMG across all trials reached 265 ± 13% of the average peak EMG in Go trials). This provides strong evidence that EMG suppression is effectively complete at this time point. On average, the EMG offset occurred significantly earlier than the SSRT at the group level (mean difference: 17 ± 6 ms, t_[16]_ = 2.59, p = 0.020, Cohen’s d = 0.63), with thirteen out of seventeen participants showing this effect (Figure 3C). Additionally, the EMG offset and SSRT were positively correlated, reinforcing their close relationship (Figure 3C).

Considering the corticospinal conduction latency of 23.3 ± 0.3 ms to the hand for these individuals (Jana et al., 2020), we estimate that cortical commands ceased around 177 ± 9 ms after the stop signal – approximately 40 ± 7 ms before SSRT. Sixteen out of seventeen individuals exhibited this pattern. Moreover, the duration of global suppression [lasting up to at least 180 ms after the stop signal, (Jana et al., 2020)], closely aligns with the timing necessary for terminating agonist EMG bursts.

These findings prompted a follow-up experiment to determine whether a similar suppression timeline applies to whole-arm reaching movements, a coordinated, multi-joint, non-ballistic action, and to examine the broader temporal dynamics of global motor suppression. In this second study, we extended the measurement window to 300 ms post-stop signal, allowing us to investigate how suppression interacts with antagonist muscle activity during movement braking and to explore suppressive mechanisms beyond discrete, ballistic tasks.

### Experiment 2

#### Stop Signal Task Performance

The probability of stopping (pStop) was ∼50% (Table 1), indicating effective implementation of the staircasing procedure. SSRT_RM_ (264 ms) fell within the typical range observed in behavioural stopping latency experiments across various effectors and devices (Brunamonti et al., 2012; Greenhouse et al., 2012; Atsma et al., 2018; Hannah et al., 2020, 2022; Jana et al., 2020; Tatz et al., 2021). Importantly, the mean SSD was ∼253 ms, which lies outside the range where violations of context independence are problematic and interfere with estimates of SSRT_RM_ (Bissett et al., 2021). Furthermore, we included SSRT_K_, a kinematic estimate of movement interruption that avoids assumptions of the Race Model, to provide a complementary and model-agnostic stopping latency measure (see below).

**Table 1.** Stop signal task performance in Experiment 2 (n = 20). Data are mean ± SEM.

| **Stop signal task indices** |  |
| --- | --- |
| Go reaction time (ms) | 508 ± 19 |
| Late stop reaction time (ms) | 465 ± 14 |
| SSD (ms) | 253 ± 23 |
| Probability of stopping (%) | 49 ± 1 |
| SSRT (ms) | 264 ± 6 |
| Go total response time (ms) | 926 ± 26 |
| **Go Errors** |  |
| Go total errors (%) | 6.3 ± 1.0 |
| **Kinematic indices** |  |
| Go peak velocity (px/s) | 3184 ± 85 |
| Late stop peak velocity (px/s) | 1826 ± 89 |
| SSRT<sub>K</sub> (ms) | 272 ± 9 |
| Trigger failures (%) | 2.8 ± 1.3 |

The average Go reaction time (Go RT) of ∼500 ms (Table 1) was within the expected range for stopping tasks (Brunamonti et al., 2012; Greenhouse et al., 2012; Atsma et al., 2018; Hannah et al., 2020, 2022; Jana et al., 2020; Tatz et al., 2021). Total response time, from Go signal to the reaching the target, averaged 926 ms, implying a movement duration of ∼400 ms. As expected, Late Stop reaction time was shorter than Go RT at the group level, and in 19/20 participants, consistent with Race Model predictions (Logan et al., 1984; Verbruggen et al., 2019). The overall Go trial error rate was acceptable (Table 1, 6.3 %; range 0.3-18.6 %), with practically all errors being movements initially directed towards the wrong target (5.9 %). Since such errors may involve suppressive mechanisms similar to stopping, these trials were excluded from analyses, including MEP comparisons.

As is common in stop-signal paradigms, participants likely engaged some degree of proactive control due to the anticipation of needing to stop. Despite this, reaching movements remained relatively fast, although not ballistic. The absence of a clear triphasic agonist–antagonist–agonist EMG pattern in Go trials (see Figures 4A and 4B), a hallmark of ballistic movements (Brown and Gilleard, 1991), supports this interpretation. This movement profile was intentional, as it better reflects naturalistic reaching behaviours. Importantly, consistent agonist EMG bursts were still observed in approximately 70% of Stop trials (see below), consistent with Experiment 1 and indicating that participants initiated a prepotent response requiring reactive stopping.

**Figure 4.**
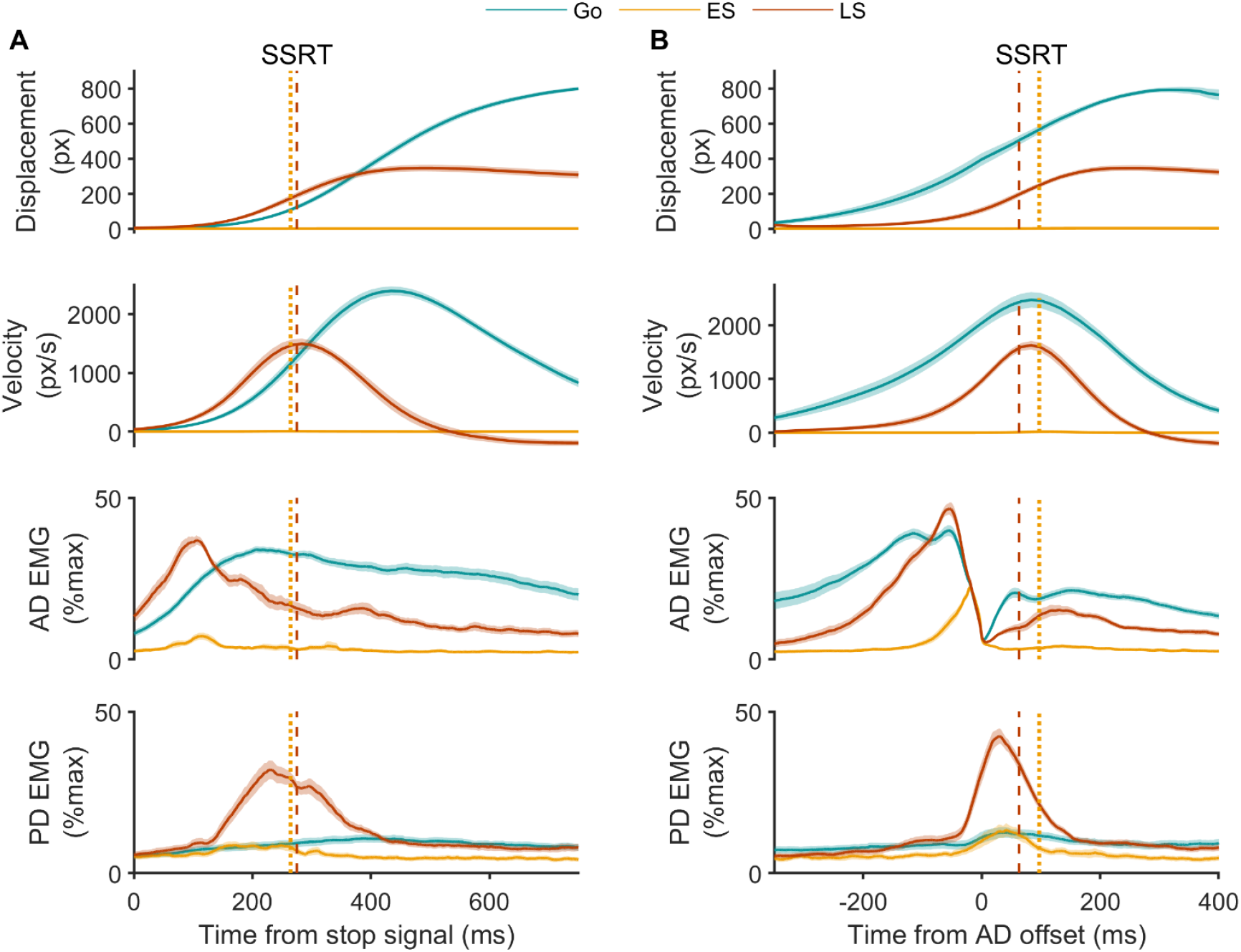
Group mean resultant displacement and velocity, and EMG from the agonist (anterior deltoid, AD) and antagonist (posterior deltoid, PD) muscles during reaching movements in Experiment 2. **A**. Data time-aligned to the stop signal on each trial, or to the most recent SSD in Go trials. Coloured lines represent different trial types: Go, Early stop (ES) and Late stop (LS). In Late Stop trials, displacement, velocity, and agonist activity are truncated relative to Go trials, indicating interruption of an already-initiated movement. This is accompanied by a burst of antagonist activity, likely reflecting active braking. In contrast, Early Stops show minimal displacement and small, brief bursts of agonist or antagonist activity. Vertical lines indicate SSRT_RM_ (dotted) and SSRT_K_ (dashed). **B**. Same data aligned time-aligned to agonist offset. Agonist activity shows a rapid decline to baseline before SSRT_RM_ (Early Stop) and SSRT_K_ (Late Stop). Antagonist activity shows a sharp increase around agonist offset in Late Stop trials, preceding SSRT_K_. Following that, both Late Stop and Go trials show a secondary agonist EMG burst, despite the absence of a stop signal in the latter, supporting the interpretation that this late activity reflects postural stabilisation of the displaced arm rather than continued movement generation or a secondary suppression. Data are group mean ± SEM.

#### Movement kinematics and muscle activity during the stop signal task

##### Go Trials: Baseline Kinematics and Muscle Activity

In Go trials, the agonist (AD) muscle exhibited a rapid increase in activation before movement onset, peaking prior to peak movement velocity (Figure 4A and 4B). Antagonist (PD) EMG activity was detected in 57 ± 7% trials. The relative infrequency and small amplitude of antagonist EMG bursts (42 ± 4% of the average peak observed in Late Stop trials), as well as their variable timing across participants, made their presence less prominent in the averaged time-series data (Figure 4A and 4B).

##### Early Stop Trials: Absent movements and truncated muscle activity

As expected, Early Stop trials showed no overt movement, with cursor displacement restricted to the ‘home’ area (Figure 4A and 4B). However, small bursts of agonist and antagonist activity were present in a subset of Early Stop trials (40 ± 5% and 47 ± 7% trials, respectively). The peak amplitude of agonist activity in Early Stops reached 17 ± 2% of the average Go trial peak, while antagonist activity reached 27 ± 4 % of the average peak observed in Late Stop trials. These low-amplitude, variably timed bursts were not apparent in the group-averaged traces time-locked to the stop signal (Figure 4A), likely due to temporal smearing (Salomoni et al., 2025). This reinforces the importance of trial-level analyses and event-aligned measures such as EMG offset to accurately capture burst dynamics (Figure 4B).

The time from stop signal to peak agonist activity, marking the onset of the decline in EMG, in Early Stops was tightly constrained (138 ± 30 ms; Figures 4A, 5, 6A). Agonist activity began to decline well before the SSRT_RM_ (Figure 4A, 4B and 5, 6A). Crucially, agonist offset occurred at 168 ± 31 ms, just 30 ± 3 ms after peak activity, and significantly earlier than the SSRT (mean difference: -97 ± 6 ms, t_[19]_ = 15.62, p = 2.7 × 10^^12^, Cohen’s d = 3.49; Figures 4B and 5, 6B). This confirms that muscle activity cessation preceded the behavioural stopping latency by a substantial margin.

Despite the lack of overt movement, antagonist bursts were recruited shortly after agonist offset at an onset of 211 ± 7 ms with respect to the stop signal (Figures 4A, 5 and 6C). Antagonist onset also significantly preceded the SSRT (mean difference: -75 ± 12 ms, t_[19]_ = 6.39, p = 3.95 × 10^^-6^, Cohen’s d = 1.42; Figures 4A, 4B, 5, 6C). Despite a trend towards a positive association between antagonist onset and SSRT in Early stops the correlation was not statistically significant (Figure 6C), potentially reflecting the lower reliability of antagonist recruitment in these trials, where movement was minimal.

##### Late Stop Trials: Interrupted Movements and Coordinated Agonist-Antagonist Dynamics

In Late Stop trials, movement was initiated but interrupted before reaching the target, as indicated by reduced peak velocity and displacement compared to Go trials (Figure 4A and 4B). Importantly, peak velocity occurred earlier in Late Stop trials compared to Go trials relative to the stop signal (Figure 4A). We interpret the timing of peak velocity in Late Stop trials as a behavioural marker of when the go-related motor command was curtailed (SSRT_K_), rather than when movement fully ceased. This better reflects the termination of descending drive, with subsequent motion continuing briefly due to mechanical inertia (see below for further discussion).

Our kinematic estimate of the latency of behavioural stopping, SSRT_K_ (272 ms, Table 1), was statistically similar to SSRT_RM_ (mean difference: 10 ± 6 ms, t_[19]_ = 1.61, p = 0.123, Cohen’s d = 0.36, BF_10_ = 0.70 in favour of null hypothesis) and the two were strongly correlated (r = 0.70, p < 0.001), suggestive of a shared underlying mechanism (Hannah et al., 2022).

Agonist activation began earlier in Late Stop trials, consistent with shorter reaction times in these trials (Figure 4A). The agonist burst to peaked earlier, had a smaller amplitude (84 ± 2% of the average Go trial maximum) and declined more rapidly than in Go trials (Figure 4A). The time from stop signal to peak agonist activity, marking the onset of agonist decline, occurred within 125 ± 45 ms (Figure 4B, 5, 6D). Agonist offset (212 ± 42 ms) followed shortly after the peak, again occurring significantly earlier than the SSRT (mean difference: -63 ± 7 ms, t_[19]_ = 9.08, p = 2.4 × 10^^-8^, Cohen’s d = 2.03; Figures 4B, 5 and 6E). Given that the corticomuscular conduction time to the AD muscle, i.e. the delay from a cortical excitation to the resulting muscle response as measured by MEP latency, is 9.6 ± 0.2 ms, the actual cessation of cortical commands must have occurred at least ∼10 ms before the observed EMG offset, and therefore well before the SSRT. Additionally, faster agonist offset was associated with shorter SSRTs (Figure 6E), reinforcing the link between early suppression at the muscle level and successful stopping.

Movement continued after the agonist burst had ended, not due to sustained motor drive, but because of mechanical inertia, a well-established feature of rapid movements (Britton et al., 1994; Kudo and Ohtsuki, 1998; MacKinnon and Rothwell, 2000). Thus, SSRT_K_ offers a practical behavioural marker of when motor output is curtailed, while subsequent deceleration and return to zero velocity reflect antagonist-mediated braking.

Antagonist bursts were present in nearly all Late Stop trials (93 ± 1%; Figures 4A,B), and typically began after agonist offset, at 195 ± 9 ms after the stop signal, but prior to SSRT_K_ (mean difference vs. SSRT_K_: -80 ± 5 ms, t_[19]_ = 19.43, p = 5.3 × 10^^-14^, Cohen’s d = 4.34; Figures 4 and 5). This temporal relationship suggests that SSRT_K_ marks a biomechanical transition: agonist-driven acceleration ceases, and antagonist-mediated braking begins. Earlier antagonist onset was also associated with shorter SSRTs (Figure 6F), further supporting its role in halting movement. These findings highlight the importance of interpreting kinematic measures like SSRT_K_ in the context of the underlying neuromuscular dynamics. While EMG offset may suggest command cessation, SSRT_K_ captures the biomechanical phase in which movement is actively slowed by reactive braking.

**Figure 5.**
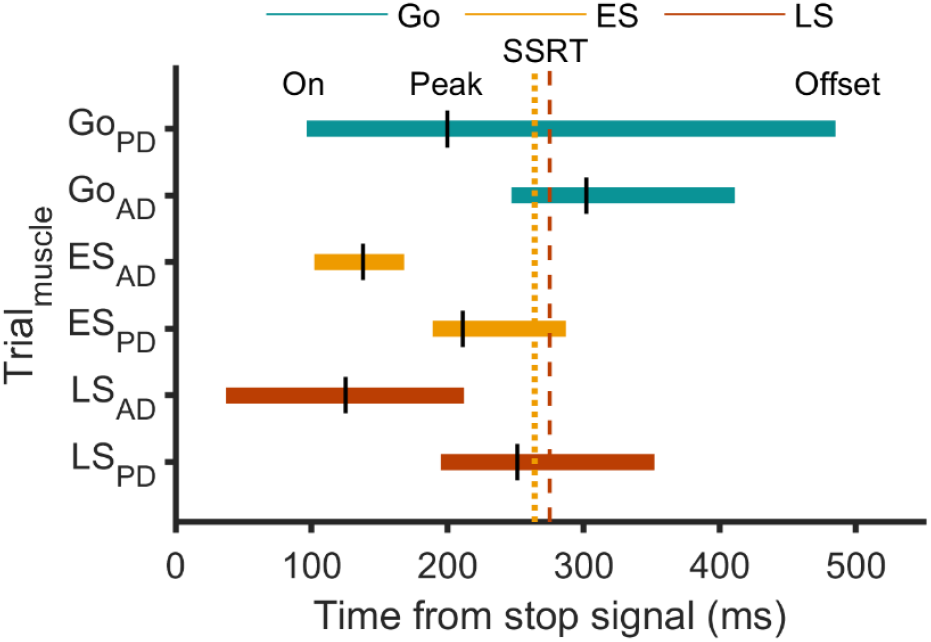
EMG burst timings with respect to the behavioural latency of stopping in Experiment 2. Timing of EMG bursts in the agonist (AD) and antagonist (PD) muscles relative to the stop signal. Coloured horizontal bars represent the mean duration of muscle activity (onset to offset) across trial types, with vertical black lines indicating mean peak timing. The vertical lines mark the mean Stop Signal Reaction Time (dotted line, SSRT_RM_, dashed line, SSRT_K_). Data are group mean.

**Figure 6.**
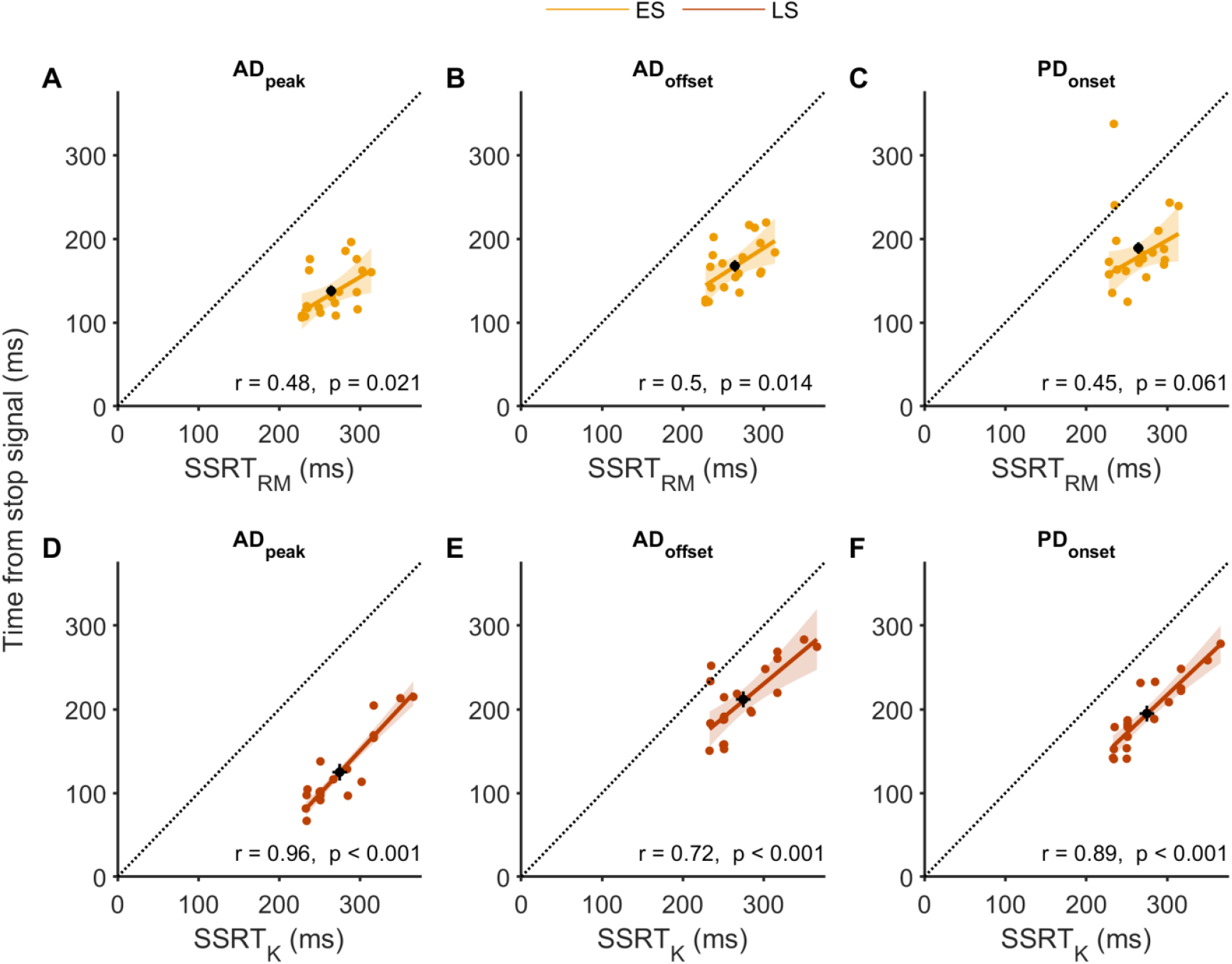
Relationships between stopping latencies (SSRT_RM_ and SSRT_K_) and EMG timings (agonist peak and offset, antagonist onset) across Early (top row, panels A–C) and Late Stop trials (bottom row, panels D–F). Each panel plots individual participant data, with the dashed black line representing unity and the black circle indicating the group mean ± SEM for SSRT and the corresponding EMG timing. In nearly all cases, data points fall below the unity line, indicating that EMG events consistently occur prior to behavioural stopping. Moderate-to-strong correlations (except in panel C) further support a systematic relationship of agonist EMG suppression and antagonist EMG activation dynamics with stopping latency.

The pattern of antagonist recruitment is consistent with prior studies of dynamic upper-limb and head movements (Kudo and Ohtsuki, 1998; Goonetilleke et al., 2010; Corneil et al., 2013; Atsma et al., 2018). In contrast, a static pressing task showed minimal antagonist activity (Van Boxtel et al., 2001), likely due to the absence of limb movement. In our data, antagonist burst amplitude scaled with the demands of stopping—larger bursts followed greater initial agonist activation (i.e. Late vs. Early Stops; Figure 4B), consistent with the need to counter greater momentum. Together, these results support the interpretation that antagonist engagement in our task reflects a reactive braking response.

Interestingly, following the initial agonist offset and antagonist onset, AD activity resumed in Late Stop trials (Figure 4B). Crucially, this secondary agonist burst does not reflect residual movement generation, but a new motor command issued after cancellation of the initial movement. In Late Stop trials, a second agonist burst began before peak velocity and persisted even after displacement and velocity had returned to zero (Figure 4B), indicating that the limb was stationary while EMG activity remained. This pattern suggests the burst supports postural stabilisation rather than continued movement.

Supporting this biomechanical interpretation, the second burst was absent in Early Stop trials, where no displacement occurred, and was more prominent in Go trials, where full reaching required greater stabilisation (Figure 4B). This graded expression according to movement extent, coupled with its persistence after movement had ceased, suggests that the activity reflects an updated postural goal rather than a continuation of the original motor plan or a second inhibitory process. Importantly, Go trials also exhibited two distinct agonist bouts of activity despite the absence of a stop signal, further confirming that the second bout is unrelated to suppressive processes and instead corresponds to postural control demands.

### Global suppression and its timing relative to EMG activity

In Go trials, MEP amplitudes in the task-irrelevant FDI muscle remained stable across all time points, except for a significant elevation relative to baseline at ∼100 ms after the stop signal (Figure 7A and 7B).

**Figure 7.**
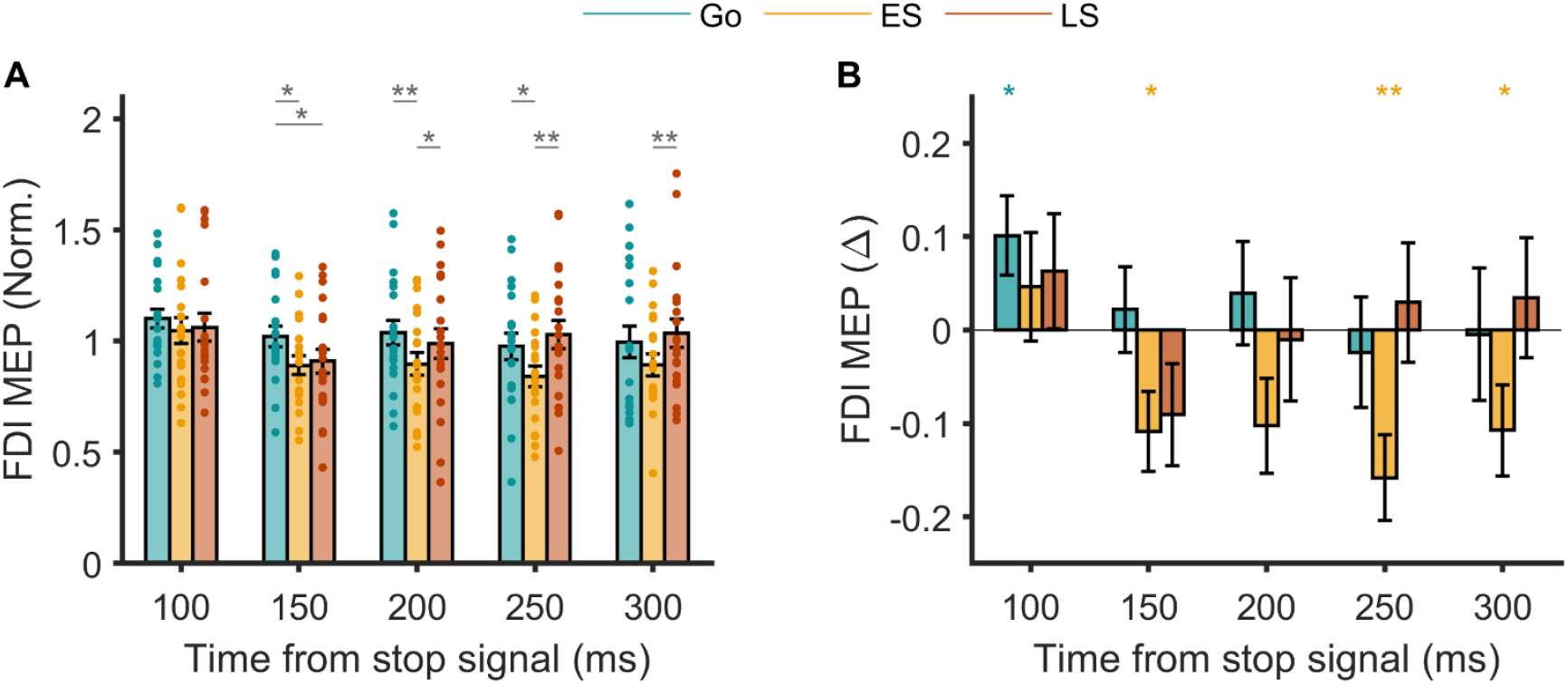
Time course of global suppression Experiment 2. **A**. MEP amplitudes across Go, Early Stop and Late Stop trials in the task-irrelevant FDI normalised to those at the Go signal (baseline). Data are group mean ± SEM with smaller symbols reflecting data from individual participants. Stars and associated horizontal bars indicate significant differences between trial types at each time point (p-value: *p, < 0.05; **, p < 0.01; ***, p < 0.001). **B**. The same data expressed as a change for each trial type relative to the baseline. Stars indicate significant differences compared to baseline for each trial type (p-value: *p, < 0.05; **, p < 0.01; ***, p < 0.001).

In Early Stop trials, MEP amplitudes were significantly reduced compared to Go trials between 150–250 ms (Figure 7A) and were notably below baseline at 150, 250 and 300 ms (Figure 7B). These findings align with prior work reporting global MEP suppression starting within 140 ms of a stop signal (Jana et al., 2020).

In Late Stop trials, MEP amplitudes briefly decreased within 150 ms, exhibiting an initial suppression similar to Early stops before returning to baseline by 200 ms (Figures 7A, 7B). This reduction was statistically significant relative to Go trials. The pattern indicates that global suppression is transiently engaged and then rapidly withdrawn as the movement is interrupted.

Repeated measures ANOVA on the normalised MEP data supported these findings, confirming a significant trial type × time interaction (F_[8,152]_ = 2.286, p = 0.024), and showing that corticospinal excitability varied across trial types over time. There was a main effect of time (F_[4,76]_ = 9.136, p = 4.42 × 10^−6^) but no effect of trial type (F_[2,38]_ = 2.21, p = 0.123).

In summary, global suppression began ∼150 ms post-stop signal, broadly aligning with the decline of agonist EMG in both Early and Late Stop trials. In Early stops, suppression persisted up to 300 ms after the stop signal, extending beyond agonist termination (∼200 ms) and SSRT (∼260 ms), overlapping with relatively weak and infrequent antagonist recruitment (∼200 ms). In Late stops, suppression was shorter-lived, returning to baseline by ∼200 ms, coinciding with agonist offset and onset of strong antagonist recruitment. These findings support a model where global suppression, agonist cancellation, and antagonist activation are temporally coordinated to either prevent or interrupt movement effectively.

## DISCUSSION

Across two experiments, we show that agonist muscle activity is fully suppressed before behavioural stopping in both ballistic and non-ballistic movements. In reaching movements, global suppression begins with agonist suppression and outlasts it when movement is cancelled early, but subsides as antagonist muscles are recruited to brake ongoing motion. These findings highlight a coordinated interplay between global suppression and agonist-antagonist dynamics during the interruption of naturalistic movements like reaching.

### Global Suppression Halts Agonist Activity

In Experiment 1, we found that agonist EMG bursts on Successful Stop trials were fully terminated ∼17 ms before the SSRT, with cortical motor commands likely ceasing even earlier (∼40 ms prior), providing clear physiological evidence that motor output is halted well before behavioural stopping. The steep decline in agonist EMG amplitude following the burst peak highlights the rapidity of this suppression. Experiment 2 extended these findings to non-ballistic, multi-joint reaching movements. Agonist bursts were abruptly truncated, whether the movement was cancelled before initiation or interrupted mid-execution. In both cases, EMG offset occurred significantly before SSRT (∼60–100 ms), further demonstrating that descending motor commands are curtailed swiftly following the stop signal. We also found that global MEP suppression began around the same time as agonist EMG suppression and persisted until complete cessation of agonist activity. This temporal alignment supports the view that global suppression is the primary mechanism responsible for cancelling motor output.

### Integration of Global Suppression and Antagonist Recruitment

The pattern of antagonist recruitment in our study highlights how action stopping integrates both neural suppression and active biomechanical braking. In Go trials, antagonist bursts were weak and inconsistent, suggesting they were not part of a strongly pre-programmed biphasic or triphasic pattern (Brown and Gilleard, 1991). In contrast, Late Stop trials exhibited prominent antagonist bursts that followed agonist offset, reflecting a reactive response to arrest ongoing movement rather than a feedforward component of the original motor plan.

Crucially, global motor suppression subsided around the time of agonist termination and the onset of these antagonist bursts. This temporal coordination supports a dynamic shift from motor output suppression to active braking. In Early Stop trials, antagonist bursts were smaller and more variable, and global suppression persisted despite their presence. This suggests that antagonist activity alone is insufficient to terminate global suppression, particularly in the absence of overt movement.

Nevertheless, antagonist activation may still support agonist suppression through reciprocal inhibition within both cortical (Griffin and Strick, 2020) and spinal cord circuits (Vallbo et al., 1979; Day et al., 1984). This additional inhibitory influence on the could contribute to a smooth transition between movement phases, reducing the need for a distinct, secondary cancellation process. Classic work shows that antagonist bursts are often preceded by agonist suppression (Hufschmidt and Hufschmidt, 1954), further supporting this possibility.

An alternative or complementary explanation is that the motor system monitors predicted versus actual sensory consequences of movement (Wolpert and Flanagan, 2001). In Late stops, ongoing movement and sensory feedback may trigger the shift from global suppression to braking. In Early Stop trials, where no movement occurs and sensory evidence is weak or lacking, global suppression may persist as a precautionary mechanism.

Although our primary focus is on cortical and corticospinal processes, this interpretation resonates with the well-established role of the cerebellum in predicting sensory outcomes (Tseng et al., 2007; Schlerf et al., 2012; Hull, 2020) and its role in initiating antagonist bursts during braking (Hallett et al., 1975; Flament and Hore, 1986). The dissociation between the timing of global suppression withdrawal and antagonist recruitment in Early versus Late Stops suggests that M1 may contribute to antagonist activation but is not essential. Subcortical structures such as the cerebellum may trigger initial antagonist bursts, with M1 providing additional modulation, consistent with its role in triphasic activation patterns typical of ballistic movements (MacKinnon and Rothwell, 2000; Irlbacher et al., 2006).

### Postural Goals and the Secondary Agonist Bout

The secondary agonist bout of activity observed in Late Stop trials is best interpreted as a postural command rather than a sign of incomplete suppression. As reported in the Results, this bout emerged after the initial reach was cancelled and persisted even after the limb came to rest, supporting a role in stabilising the arm. Its presence in Go trials and absence in Early Stops further suggest it is unrelated to stopping per se and instead reflects biomechanical demands.

These observations, combined with the timing of global suppression and antagonist braking, point to a dynamic sequence of control: global suppression halts the initial command, antagonists brake the movement, and a new postural goal is issued. In this sense, Late stopping may resemble a stop-then-update process, where the motor system shifts rapidly from cancelling the initial action to initiating a stabilising response tailored to the current biomechanical state. More broadly, these findings reveal a coordinated interplay between suppressive and facilitatory mechanisms in action stopping and updating.

While our task involved instructed stopping, many real-world scenarios require redirection rather than full cancellation. Future work should explore how global suppression operates during flexible goal updating, to better define the boundaries of its role in adaptive motor control.

### Implications for Theories and Measures of Action Stopping

The “pause-then-cancel” model posits that a secondary cancellation process follows an initial pause (Schmidt and Berke, 2017; Diesburg and Wessel, 2021). One argument in support of this model is that while EMG suppression begins before the SSRT, full termination of EMG activity takes much longer, suggesting a second process may be required to complete the stop (Hervault et al., 2025). However, our findings challenge this interpretation. By aligning EMG to its point of offset, we show that termination occurs rapidly and well before SSRT in both button-press and reaching movements. This suggests that a single, rapidly engaged suppressive mechanism can account for movement cancellation without invoking a distinct later-stage process.

While we cannot exclude the possibility that additional inhibitory mechanisms contribute to stopping, particularly within basal ganglia circuits, their dynamics in humans remain insufficiently characterised. Rodent studies have shown very early STN activity following stop signals (Schmidt et al., 2013; Mallet et al., 2016), which has been interpreted as evidence for an initial “pause” signal (Schmidt and Berke, 2017). However, in humans and non-human primates, STN activity typically emerges closer to the time of behavioural stopping and differentiates between successful and failed stops (Bastin et al., 2014; Benis et al., 2016; Wessel et al., 2016; Pasquereau and Turner, 2017), a pattern that contrasts with rodent findings. These species differences raise important questions about the generalisability of the two-stage pause-then-cancel model. In the absence of direct human evidence for a subcortical “cancel” process, we interpret the rapid, complete suppression of EMG, closely aligned with global suppression, as a sufficient physiological marker of stopping. Further work will be needed to clarify whether pallido-striatal activity in humans plays an additional, functionally distinct role beyond the STN-mediated suppression already evident at cortical and muscular levels.

Our finding that agonist activity is terminated before SSRT has implications for putative indices of the stop process. The P3 EEG marker (Wessel and Aron, 2015; Huster et al., 2020; Hervault et al., 2025), has been associated with the ‘cancel’ process in the two-stage model, since it appears to occur around the SSRT (Hervault et al., 2025). Although we did not measure EEG in the present study, our observation that muscle activity is already fully suppressed prior to SSRT raises a testable question about whether additional cancellation mechanisms are still necessary at this stage. One possibility is that the P3 reflects a downstream monitoring or confirmation process, rather than an active cancellation step per se (Huster et al., 2020; Hervault et al., 2025). This hypothesis should be directly addressed in future studies using simultaneous EEG and EMG, ideally alongside measures of subcortical activity, to clarify the functional role of the P3 in relation to the full suppression of muscle output.

More broadly, our findings suggest that global suppression is not a rigid, top-down mechanism but instead interacts with antagonist recruitment to flexibly regulate movement termination. This supports the idea that movement suppression and initiation are not separate processes but part of a continuous, dynamic control system. The timing of global suppression relative to antagonist recruitment suggests that action stopping is embedded within a broader framework of action selection and updating (Mostofsky and Simmonds, 2008; Du et al., 2024), aligning with views that the right inferior frontal gyrus-driven “action stopping” network is involved in action updating rather than pure behavioural suppression (Buch et al., 2010; Verbruggen et al., 2010).

## CONCLUSIONS

Our findings challenge the pause-then-cancel model of action stopping by showing that agonist activity is abruptly terminated in synchrony with global motor suppression and before behavioural stopping, eliminating the need for a second process. Additionally, global motor suppression is dynamically coordinated with antagonist recruitment rather than acting as a rigid mechanism, highlighting its integration within broader motor control. This underscores the need to refine action stopping models.

## ACKNOWLEDGEMENTS

We are grateful to John Rothwell, Sumitash Jana and Vignesh Muralidharan for their thoughtful comments on an earlier version of the manuscript.

## CODE AVAILABILITY

Data and code are available upon request.

